# Immunogenomic profile of colorectal cancer response to immune checkpoint blockade

**DOI:** 10.1101/2020.12.15.422831

**Authors:** Michele Bortolomeazzi, Mohamed Reda Keddar, Lucia Montorsi, Amelia Acha-Sagredo, Lorena Benedetti, Damjan Temelkovski, Subin Choi, Nedyalko Petrov, Katrina Todd, Patty Wai, Jonny Kohl, Tamara Denner, Emma Nye, Robert Goldstone, Sophia Ward, Gareth A. Wilson, Maise Al Bakir, Charles Swanton, Susan John, James Miles, Banafshe Larijani, Victoria Kunene, Elisa Fontana, Hendrik-Tobias Arkenau, Peter J. Parker, Manuel Rodriguez-Justo, Kai-Keen Shiu, Jo Spencer, Francesca D. Ciccarelli

## Abstract

Colorectal cancers (CRCs) show variable response to immune checkpoint blockade, which can only partially be explained by the variability of tumour mutational burden. To dissect the cellular and molecular determinants of response we performed a multi-omic screen of 721 cancer regions from patients treated with Pembrolizumab (KEYNOTE 177 clinical trial) or Nivolumab. Multi-regional whole exome, RNA and T-cell receptor sequencing show that, within hypermutated CRCs, response to both anti-PD1 agents is not positively associated with tumour mutational burden but with high clonality of immunogenic mutations, expanded T cells, low activation of the WNT pathway and active immune escape mechanisms. Coupling high-dimensional imaging mass cytometry with multiplexed immunofluorescence and computational spatial analysis, we observe that responsive hypermutated CRCs are rich in cytotoxic and proliferating PD1-expressing CD8 cells interacting with high-density clusters of PDL1-expressing antigen presenting macrophages. We propose that anti-PD1 agents release the PD1-PDL1 interaction between CD8 T cells and macrophages thus promoting cytotoxic anti-tumour activity.

## INTRODUCTION

Anti-cancer therapy based on immune checkpoint blockade has driven a paradigm shift in the treatment of several cancer types^1^. Pembrolizumab and Nivolumab, two antibodies targeting programmed cell death 1 (PD1) expressed on T cells, have shown efficacy in advanced hypermutated colorectal cancers (CRCs)^2^. Response is thought to depend on rich immune infiltration^3^ and high tumour mutation burden (TMB) leading to increased production of peptide neoantigens. However, despite pervasive tumour immunogenicity, response is highly variable and approximately half of patients with hypermutated CRCs show no durable benefit^4^.

We have dissected the extent to which TMB, cancer dysfunctional genes and pathways as well as the qualitative and quantitative immune composition of the tumour microenvironment (TME) influence response to immune checkpoint blockade. To reproduce the most common clinical scenario where metastatic biopsies are not routinely taken, we performed a high-dimensional and multi-regional profile of primary CRCs or local relapses from 29 patients, divided into a discovery and a validation cohort. The discovery cohort was composed of patients subsequently treated with Pembrolizumab as first-line therapy within the KEYNOTE 177 phase III clinical trial^5^ or Nivolumab in advanced metastatic setting. The fact that most patients did not receive previous treatment offered the ideal opportunity to identify critical factors for response to treatment in terms of cancer genetic and transcriptional dysregulation and immune microenvironment composition. We then extended the study to a validation cohort of patients who received anti-PD1 agents in chemorefractory setting to assess the general validity of our findings.

## RESULTS

### Response of hypermutated CRCs is associated with clonal immunogenic mutations and clonally expanded T cells

To assess how immune infiltration correlates with tumour genetic and transcriptional alterations in CRC, we performed a multi-omic and multi-regional profile of 24 sequential slides (A-K) from formalin-fixed paraffin-embedded (FFPE) tumour blocks, for a total of 555 regions from 16 patients of the discovery cohort (Fig.1A). Ten of these patients received Pembrolizumab (UH1-UH10) and six Nivolumab (UH11-UH16) in advanced metastatic setting. Overall, nine patients achieved durable benefit and seven had no durable benefit from the treatment according to RECIST 1.1 (Supplementary Table 1). We validated the main findings of the study in 166 additional regions from 13 CRC patients (UH17-UH29, Extended Data Fig.1A) treated with anti-PD1 agents alone or in combinations with other immune checkpoint inhibitors (Supplementary Table 1). Ten of them reached a durable benefit and three had no durable benefit from the treatment (Fig.1A).

**Fig. 1.**
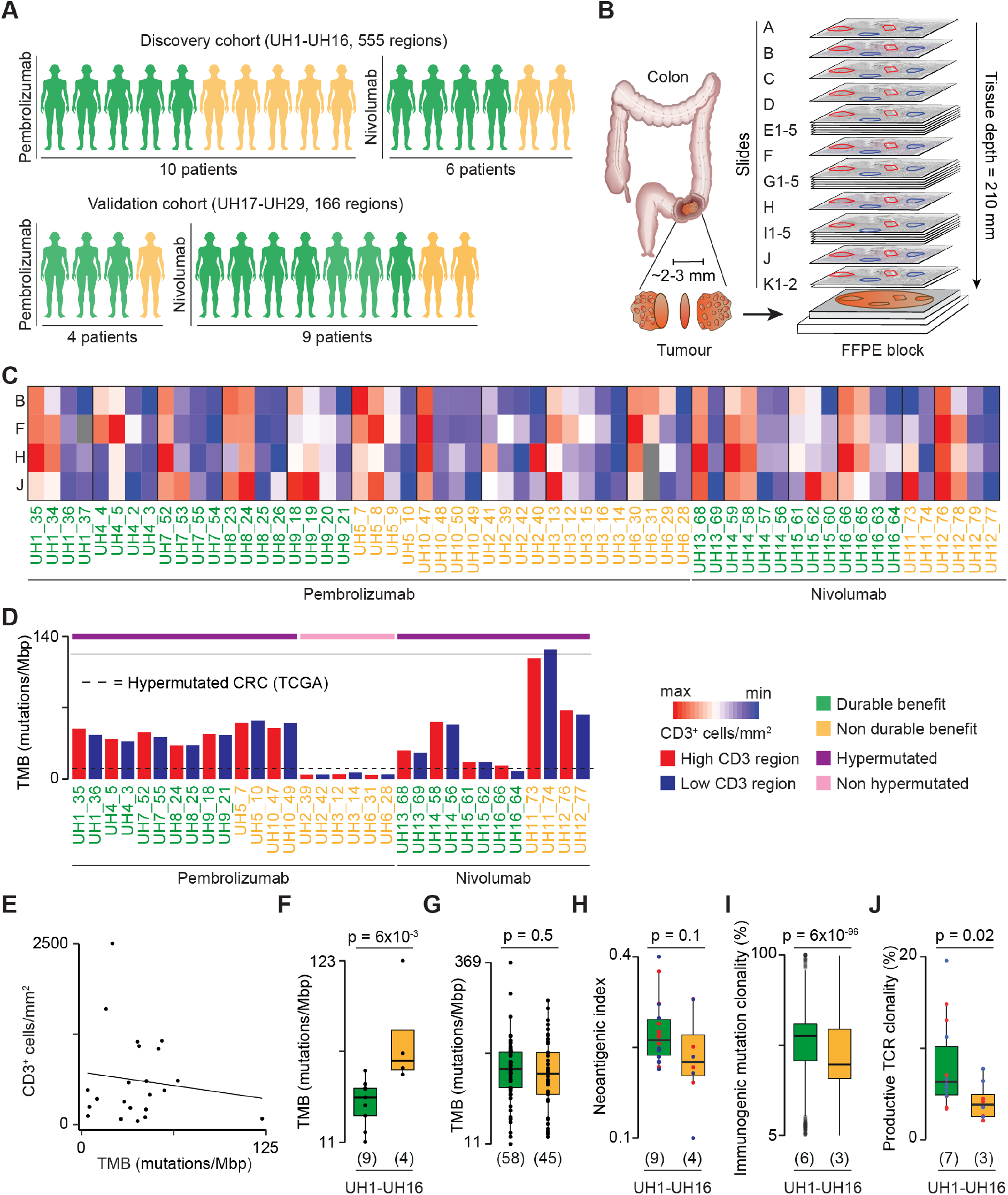
Study design and quantification of tumour heterogeneity. **(A)** Description of the study cohorts. Clinical benefit from the treatment was assessed with RECIST 1.1. **(B)** Experimental design for the discovery cohort. Twenty-four sequential slides from FFPE CRC blocks prior to treatment were used for multiregional CD3 immunohistochemistry (slides A, B, F, H and J), IMC (slide C), mIF (slide D), WES (slides E1-5), RNA-seq (slides G1-5), TCR-seq (slides I1-5) and A-FRET detection of PD1-PDL1 interaction *in situ* (slides K1-2). Multiple regions with variable CD3 infiltration were identified in slide A and projected to all other slides. **(C)** Quantification of CD3^+^ cells/mm^2^ from immunohistochemistry staining in 60 regions of the discovery cohort using Qupath^22^. Values were normalised within each patient. Grey boxes indicate missing measures. **(D)** TMB of 32 sequenced regions in the discovery cohort. The dotted line corresponds to the TMB of hypermutated CRC (12 mutations/Mbp)^6^. **(E)** Correlation between CD3^+^ cells/mm^2^ from immunohistochemistry staining of slide F (discovery) and E (validation) and TMB across samples. Average CD3^+^ cell density across multiple regions per slide is reported. For the discovery cohort, TMB was calculated as the average between the two sequenced regions. For the validation cohort, TMB was obtained from the FM1 test^31^. Comparison of TMB between DB- and nDB-CRCs of the discovery cohort **(F)** and in an extended cohort of hypermutated samples from the validation cohort (Supplementary Table 1) and published studies ^7–11^ **(G)**. For^9,11^ response was unavailable and overall survival from the start of immunotherapy was used to define DB (≥ 12 months) and nDB (< 12 months). Comparison of **(H)** ratio of predicted immunogenic mutations (SNVs and indels) over all nonsilent mutations (neoantigenic index); **(I)** clonality of immunogenic mutations in 22 regions with >30% purity (Supplementary Table 2). Regions with lower purity were excluded because of unreliable mutation clonality assessment; and **(J)** productive clonality of T-Cell Receptor (TCR) beta rearrangements between DB- and nDB-CRCs. Number of patients in each tumour group is reported in brackets. Distributions were compared using two-sided Wilcoxon test.

Through pathological inspection of CD3 immunostaining, we identified multiple regions per block with variable T cell content in proximity to the tumour infiltrating margins (Extended Data Fig.1B). These regions were then projected in all sequential slides to perform additional CD3 immunohistochemistry, imaging mass cytometry (IMC), multiplexed immunofluorescence (mIF), whole exome sequencing (WES), RNA sequencing (RNA-seq), T-cell receptor β-chain sequencing (TCR-seq) and amplified Förster Resonance Energy Transfer (A-FRET) detection of PD1-PDL1 interaction *in situ* (Fig.1B).

T cells are the effector cells that mediate the response to anti-PD1 immunotherapy. Therefore, we compared T cell infiltration between and within tumours (Extended Data Fig.1C). In both discovery (Fig.1C) and validation (Extended Data Fig.1D) cohorts, we observed widespread inter- and intra-tumour heterogeneity of T cell infiltration with up to 38-fold difference in CD3 cell densities between patients and up to 20-fold difference between regions of the same patient (Supplementary Table 2). To investigate how heterogeneity in T cell infiltration correlated with TMB, we performed multiregional WES in the discovery cohort selecting two regions per patient, one with high and one with low T cell infiltration (Supplementary Table 2). TMB was comparable between regions of the same patient (Fig.1D) and did not correlate with T cell content across samples (Fig.1E), indicating its independence of inter- and intra-tumour T cell heterogeneity.

WES analysis also showed that three patients (UH2, UH3, UH6) had a TMB lower than 12 mutations/Mbp (TCGA lower bound of CRC hypermutated phenotype^6^) despite negative MLH1 and PMS2 immunostaining and consistent with resistance to treatment (Supplementary Table 1). All patients of the validation cohort had hypermutated CRCs (Supplementary Table 1).

Given that approximately 50% of patients with hypermutated CRC do not respond to immunotherapy, we compared TMB between hypermutated CRCs with durable benefit (DB-CRCs) and those with no durable benefit (nDB-CRCs) to assess the role of TMB as a marker of response. Surprisingly, in the discovery cohort DB-CRCs had a significantly lower TMB than nDB-CRCs (Fig.1F). When adding further hypermutated CRCs from the validation cohort and published studies^7^-^11^, we observed no significant difference between DB- and nDB-CRCs (Fig.1G). Therefore, in CRC a TMB below 12 mutations/Mbp is a predictor of resistance to anti-PD1 immunotherapy while, above this threshold, it is no longer associated with response.

To understand whether the proportion of cancer-associated neoantigens differed between responders and non-responders, we predicted how many cancer mutations were potentially immunogenic in each sample. In the discovery cohort, the ratio between immunogenic mutations and all mutations (neoantigenic index) was similar between DB-CRCs and nDB-CRCs (Fig.1H). However, immunogenic mutations were more clonal in DB-CRCs (Fig.1I) indicating expansion of tumour cells displaying the same potential immune targets. Consistent with dominant antigenic targets, the productive TCR repertoire was also more clonal in DB-CRCs (Fig.1J).

Therefore, hypermutated CRCs responding to anti-PD1 agents have significantly more clonally expanded immunogenic mutations and T cells compared to those failing to respond.

### DB-CRCs show widespread immune dysregulation and silencing of the *B2M* gene

To dissect CRC molecular determinants of response to anti-PD1 agents, we compared genetic and transcriptional dysregulations between hypermutated and non-hypermutated CRC as well as between DB-CRCs and nDB-CRCs.

Genes of the WNT pathway were frequently damaged (Extended Data Fig.2A, Supplementary Tables 3) and transcriptionally deregulated (Fig.2A, Supplementary Tables 4) in hypermutated compared to non-hypermutated CRCs. Moreover, the downstream targets of WNT were significantly downregulated in hypermutated compared CRCs in our cohort (Fig.2B) as well as in the whole TCGA (Fig.2C). Since transcriptional WNT activation is known to reduce T cell infiltration^12^,^13^, this suggests a potential impact on the TME of non-hypermutated CRCs.

**Fig. 2.**
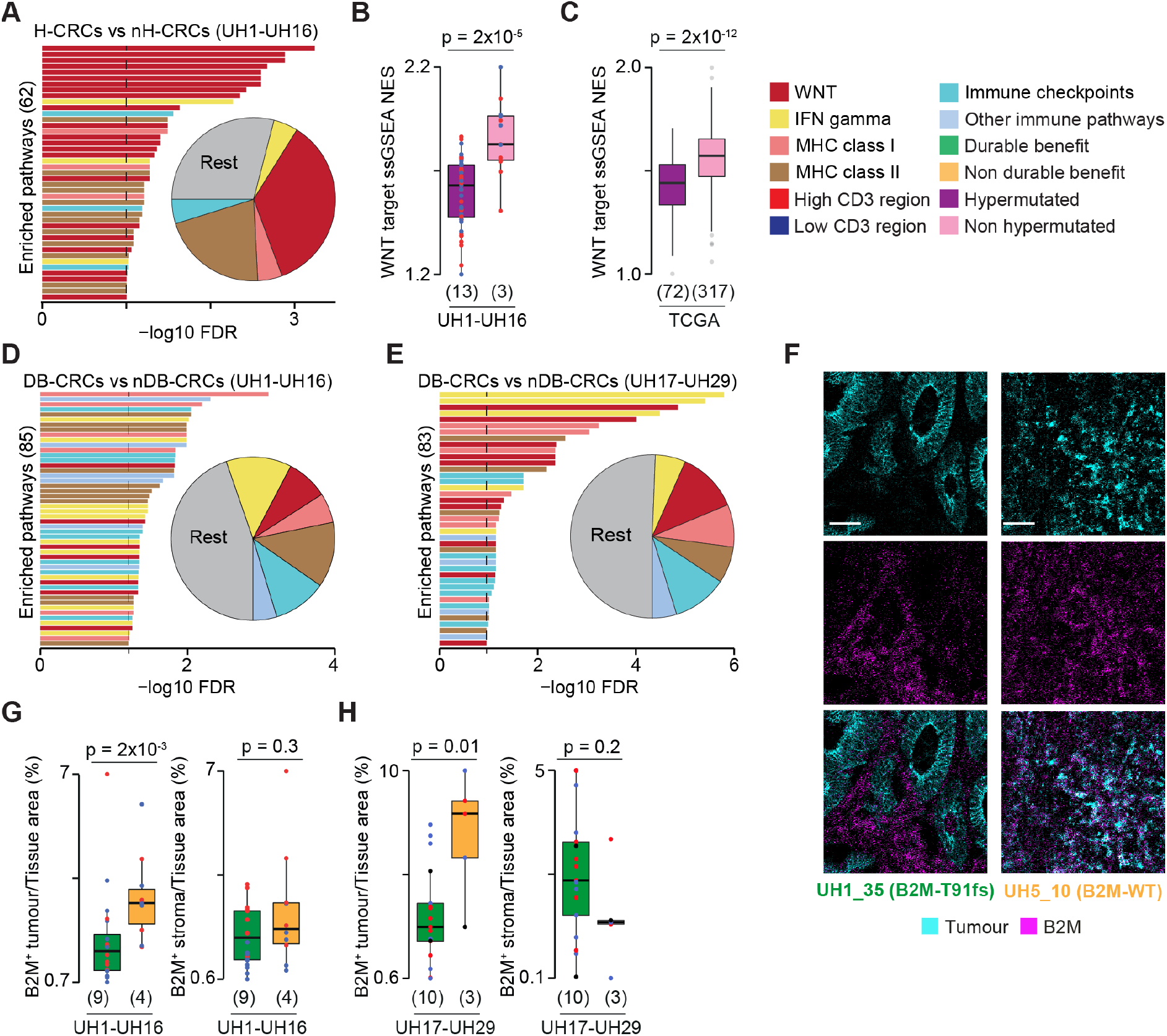
Cancer driver and immune aberrations across CRC groups. **(A)** Representative pathways enriched in differentially expressed genes (DEGs) between hypermutated and non-hypermutated CRCs of the discovery cohort. Normalised enrichment scores (NES) from Single sample Gene Set Enrichment Analysis (ssGSEA)^48^ of 68 transcriptional targets of the WNT pathway^12^,^49^ between hypermutated and non-hypermutated CRCs from the discovery cohort (**B**) and The Cancer Genome Atlas (TCGA) (**C**). Representative pathways enriched in DEGs between DB and nDB CRCs from the discovery **(D)** and validation **(E)** cohorts. In panels (**A,D,E**) False Discovery Rate (FDR) was calculated using Benjamini-Hochberg correction (10% for pathway enrichment and 5% for DEGs). Proportions of immune-related pathways over all enriched pathways are reported as pie charts. **(F)** Representative IMC images of CRCs with mutated and wild-type (WT) B2M protein. Scalebar = 50 μm Comparison of normalised tumour and stroma B2M^+^ areas between DB- and nDB-CRCs of the discovery (**G**) and validation (**H**) cohorts. Number of patients in each tumour group is reported in brackets. Distributions were compared using two-sided Wilcoxon test. IFN, Interferon; MHC, Major histocompatibility complex.

Genes encoding members of the interferon gamma pathway, antigen presentation machinery and other immune-related processes were damaged or transcriptionally dysregulated in DB-compared to nDB-CRCs in the discovery (Fig.2D Supplementary Table 4) and validation (Fig.2E) cohorts. Interestingly, two DB-CRCs of the discovery cohort showed a clonal truncating mutation (T91fs) in the beta-2-microglobulin (B2M) gene encoding the invariable subunit of the MHC class I complex (Extended Data Fig.2B). Since a B2M antibody was part of the IMC panel (Supplementary Table 5), we could assess that *B2M* truncating mutation resulted in no protein expression in the tumour, as compared to a widespread B2M expression in patients with wild-type *B2M* (Fig.2F). In general, B2M protein expression was significantly reduced in the tumour but not in the stroma of DB-CRCs in the discovery (Fig.2G) and validation (Fig.2H) cohorts as well as in both cohorts combined (Extended Data Fig.2C). In melanoma, *B2M* loss has been associated with resistance to immune checkpoint inhibitors^14^. Our data indicate an opposite association in CRC, supporting similar recent observations^15^.

### Hypermutated CRCs are enriched in cytotoxic and proliferating CD8 T cells

To understand the role of TME in the response to anti-PD1 agents, we analysed multiple tumour regions of the discovery cohort with IMC (Extended Data Fig.3A). We used thirty markers identifying T cells, macrophages, neutrophils, dendritic cells, B cells as well as the tumour structure (Supplementary Table 5). After regional ablation and image processing, we derived tumour and stroma masks for each region (Fig.3A) and verified that the relative proportion of stroma and tumour cells was similar across samples (Extended Data Fig.3B-F). We then analysed the TME images through two independent approaches. In one, we measured the normalised pixel area of individual or combined markers (pixel analysis, Fig.3A). In the other, we performed single cell segmentation, assigned cell identities and measured the relative abundance of immune cell populations identified through unsupervised single cell clustering (single cell analysis, Fig.3A). Outcomes of all computational analyses were validated by independent manual assessment of unprocessed images.

**Fig. 3.**
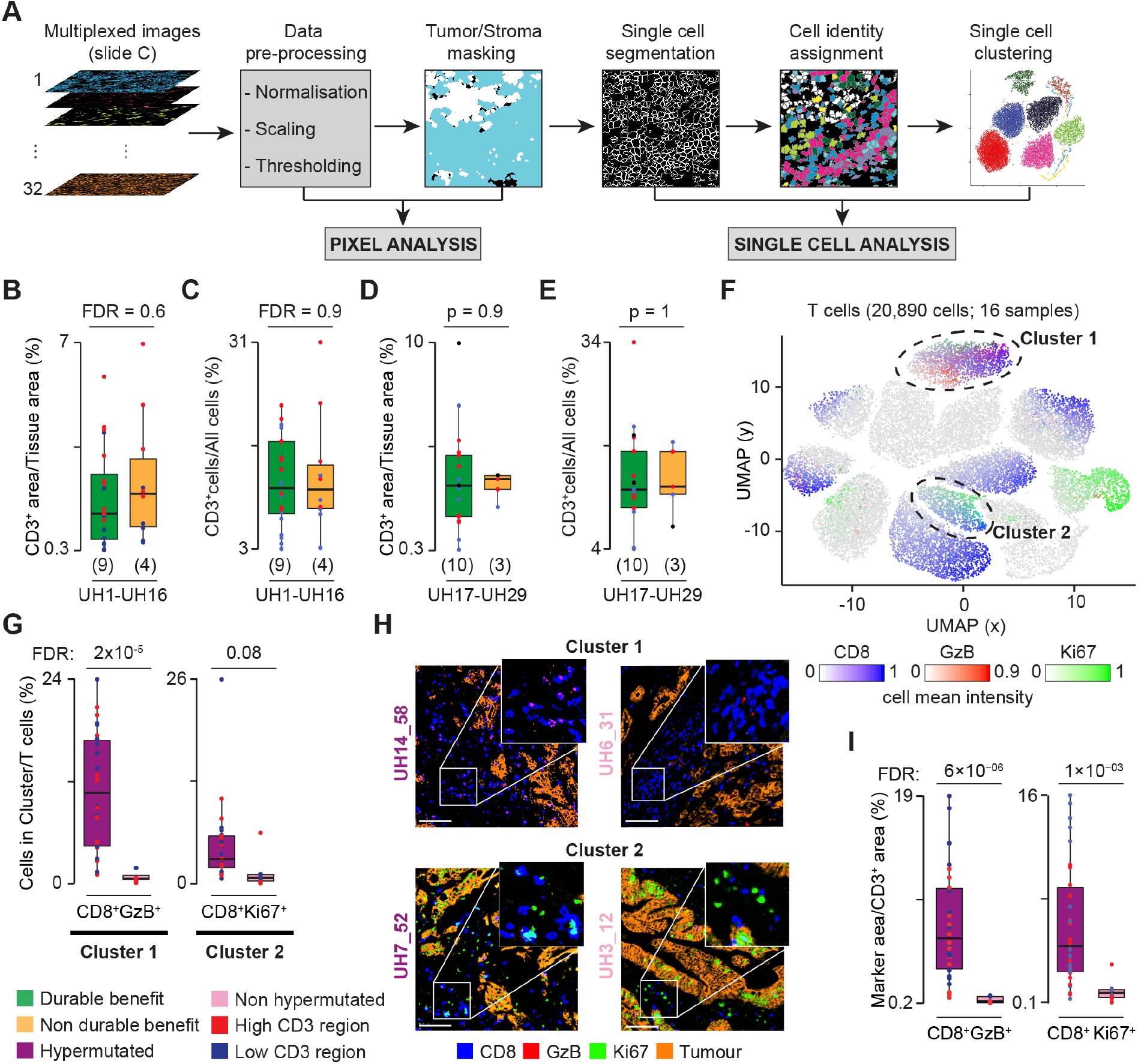
Comparison of T cells infiltrates between CRC groups. **(A)** Imaging Mass Cytometry (IMC) analytical workflow. For each region, images representing the marker expression patterns (Supplementary Table 5) and DNA intercalators were pre-processed to extract pixel intensities. Masks for tumour and stroma were derived and used for the pixel analysis. Each region was segmented into single cells that were assigned to tumour or stroma, phenotypically identified through expression of representative markers and used for single cell clustering. Comparison of normalised CD3^+^ areas (**B**) and CD3^+^ cells (**C**) between DB and nDB-CRCs in the discovery cohort. Benjamini-Hochberg FDR correction was applied to account for testing over five immune populations. Comparison of normalised CD3^+^ areas (**D**) and CD3^+^ cells (**E**) between DB and nDB-CRCs in the validation cohort. No correction for multiple test was applied because only T cells were compared. **(F)** Uniform Manifold Approximation and Projection (UMAP) map of 20,890 T cells in 38 regions from 16 CRCs of the discovery cohort. Cells were grouped in 13 clusters based on the expression of 12 phenotypic markers using Seurat^26^ (Supplementary Table 7). Cells are coloured according to GzB, Ki67 and CD8 mean intensities and the two clusters enriched in hypermutated CRCs are circled. **(G)** Proportions of cluster 1 (CD8^+^GzB^+^ cells) and cluster 2 (CD8^+^Ki67^+^ cells) over the total T cells in hypermutated and non-hypermutated CRCs. Distributions were compared using two-sided Wilcoxon test. Benjamini-Hochberg FDR correction was applied for testing over 13 clusters. **(H)** IMC-derived images of tumour-associated markers (E-cadherin and Pan-Keratin) and CD8 and GzB or CD8 and Ki67 in two representative hypermutated and non-hypermutated CRCs. Scale bar = 100μm. **(I)** Comparisons of normalised CD8^+^GzB^+^ and CD8^+^Ki67^+^ areas between hypermutated and non-hypermutated CRCs. Distributions were compared using two-sided Wilcoxon test. Benjamini-Hochberg FDR correction was applied to account for testing over 25 combinations of T cell markers (Supplementary Table 6).

Hypermutated DB- and nDB-CRCs showed no difference in the normalised CD3^+^ area (Fig.3B, Supplementary Table 6) and proportion of CD3^+^ cells (Fig.3C, Supplementary Table 8), confirming that overall T cell infiltration does not correlate with TMB (Fig.1E) or response to therapy in this group. To further investigate whether DB- and nDB-CRCs differed in specific T cell subpopulations, we performed single cell clustering using 12 phenotypic markers (Supplementary Table 5). We found no qualitative or quantitative differences in T cell subpopulations between DB and nDB-CRCs (Supplementary Table 7, Extended Data Fig.4). Given their relevance to immune checkpoint inhibitors, we further profiled T cells in the validation cohort adding five markers of T cell function to the 12 used in the discovery cohort (Supplementary Table 5). We confirmed no significant difference in the normalised CD3^+^ area or proportion of CD3^+^ cells between DB- and nDB-CRCs of the validation cohort (Fig.3D,E). Moreover, when performing single cell clustering with all 17 phenotypic markers of T cells, we observed no difference in T cell infiltrates between DB- and nDB-CRCs (Supplementary Table 7).

We repeated the same comparison of T cell infiltrates between hypermutated and non-hypermutated CRCs of the discovery cohort. In this case, we found that two clusters of CD8 cells (cluster 1, expressing Granzyme B, GzB, and cluster 2, expressing Ki67) were significantly more abundant in hypermutated CRCs (Fig.3 F-H). Pixel analysis confirmed these results (Fig.3I).

Our analysis shows substantial differences in T cell infiltration between hypermutated and non-hypermutated CRCs, identifying the specific populations responsible for this difference. Non-hypermutated CRCs are highly depleted of cytotoxic and proliferating CD8 T cells, consistent with the activation of the WNT pathway (Fig.2A,B). No qualitative or quantitative differences in T cells were detected between hypermutated DB and nDB-CRCs, which are both rich in CD8 T cells.

### Hypermutated DB-CRCs are enriched in CD74^+^ macrophages

We undertook single cell clustering of all other main immune populations (macrophages, dendritic cells, neutrophils and B cells) in the discovery cohort using sets of phenotypic markers specific for each of them (Supplementary Table 5). Applying the same strategy used for T cell analysis, we compared the relative abundance of the obtained cell clusters between hypermutated and non-hypermutated CRCs or DB and nDB-CRCs.

We found no difference in any other immune cell population between hypermutated and non-hypermutated CRCs (Supplementary Table 7). However, we observed proportionally higher CD68^+^CD74^+^ cells in DB-CRCs compared to nDB-CRCs (cluster 3, Fig.4A-C), which was confirmed also by pixel analysis (Fig.4D). To validate these results, we profiled the macrophages in the validation samples using the same phenotypic markers of the discovery cohort. Pixel analysis confirmed higher normalised CD68^+^CD74^+^ area in the validation samples alone (Fig.4E) and when all hypermutated CRCs were analysed together (Fig.4F). To identify CD68^+^CD74^+^ cells, we applied a threshold of 0.1 CD74 expression to all macrophages in the validation cohort (Fig.4G) and in all hypermutated CRCs (Fig.4H). We verified that CD68^+^CD74^+^ cells identified in this way matched phenotypically to cells in cluster 3 of the discovery cohort (Extended Data Fig.5A-C). We then compared the proportion of CD68^+^CD74^+^ cells between DB-CRCs and nDB-CRCs and found that it was higher in DB-CRCs of the validation cohort alone (Fig.4I) and when all hypermutated CRCs were analysed together (Fig.4J).

**Fig. 4.**
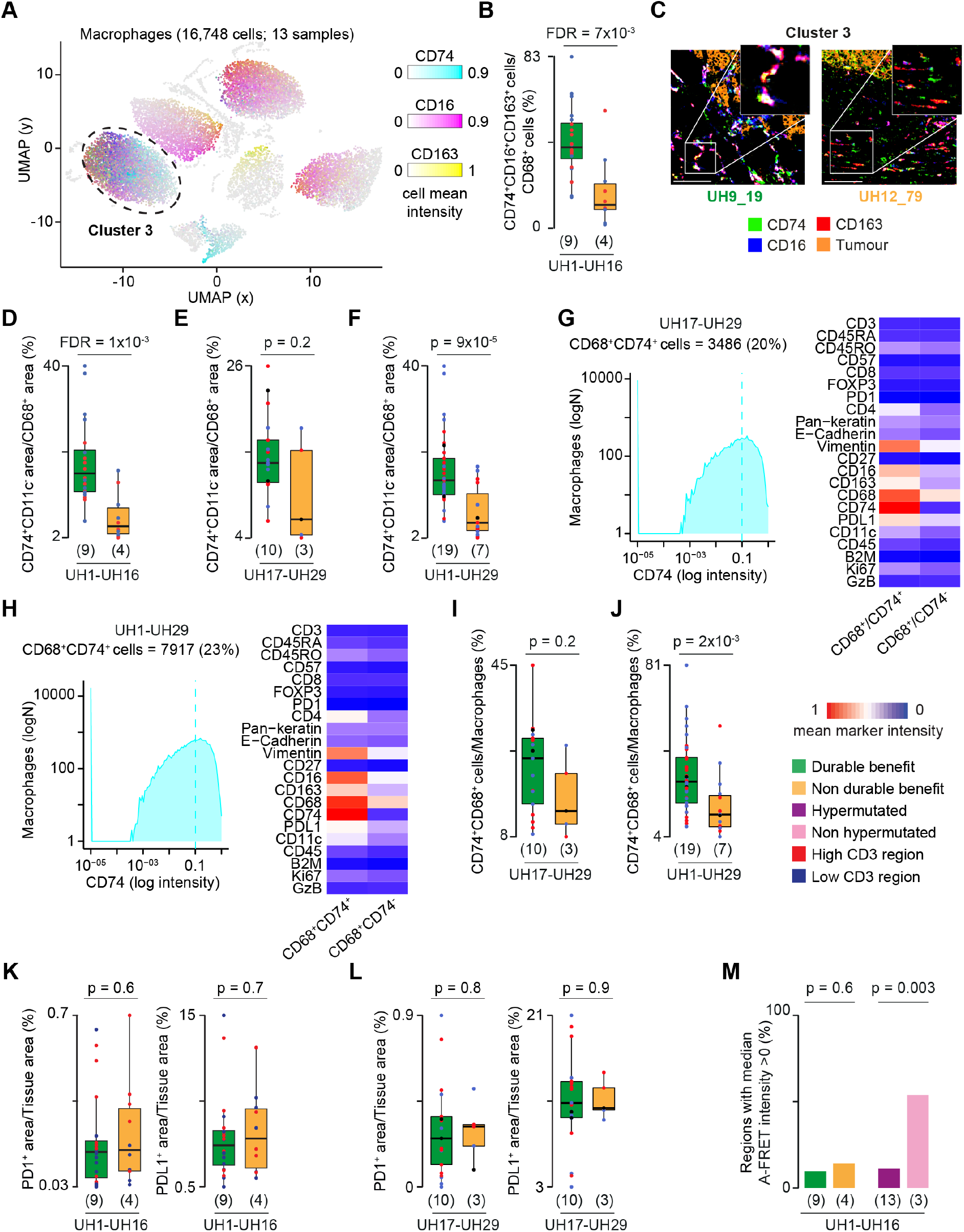
Difference in CD74^+^ macrophages between DB and nDB-CRCs. **(A)** UMAP map of 16,748 macrophages in 30 regions from 13 hypermutated CRCs in the discovery cohort. Cells were grouped in nine clusters based on the expression of 11 phenotypic markers using Seurat^26^ (Supplementary Table 7). Cells are coloured according to CD74, CD16 and CD163 mean intensities. **(B)**Proportions of cluster 3 (CD74^+^ cells) over the total macrophages in DB and nDB-CRCs. Distributions were compared using two-sided Wilcoxon test. Benjamini-Hochberg FDR correction was applied for testing over nine clusters. **(C)** IMC-derived images of CD74, CD16 and CD163 and tumour-associated markers (E-cadherin and Pan-Keratin) in two representative DB and nDB-CRCs. Scale bar = 100μm. Comparisons of normalised CD74^+^ area between DB and nDB-CRCs in the discovery (**D**), validation (**E**) and combined (**F**) cohorts using two-sided Wilcoxon test. For the discovery cohort, Benjamini-Hochberg FDR correction was applied for the nine tested combinations of macrophage markers (Supplementary Table 6). CD74^+^ macrophages in the validation cohort (**G**) and in all samples together (**H**) were identified by applying a threshold of 0.1 CD74 expression to all macrophages. This threshold was identified through IMC image histological inspection. The mean intensities of IMC markers in CD74^+^ and CD74^-^ macrophages are reported. Colour gradient was normalised across all markers and cells. Comparison of normalised of CD74^+^ macrophages between DB and nDB-CRCs of the validation **(I)** and combined **(J)** cohorts. Distributions were compared using two-sided Wilcoxon test. Comparison of normalised PD1^+^ and PDL1^+^ areas between DB and nDB-CRCs from the discovery (**K**) and validation (**L**) cohorts. Distributions were compared using twosided Wilcoxon test. **(M)** Percentage of regions with median amplified-Förster Resonance Energy Transfer (A-FRET) intensity higher than zero in DB and nDB-CRCs and hypermutated and nonhypermutated CRCs. Proportions were compared using Fisher’s exact test. Number of patients in each tumour group is reported in brackets.

Finally, given their role in anti-PD1 immunotherapy, we compared the normalised PD1 or PDL1 levels between DB- and nDB-CRCs and found no significant differences in the discovery (Fig.4K), validation (Fig.4L) and combined (Extended Data Fig.5D) cohorts. This was supported by gene expression analysis (Extended Data Fig.5E,F) and single cell clustering, which detected no qualitative or quantitative differences in PD1^+^ or PDL1^+^ cells between DB- and nDB-CRCs (Supplementary Table 7). In general, we observed low expression of both genes in all CRCs in our study (Extended Data Fig.5G) as well as in TCGA, where *PD1* and *PDL1* gene expression in CRC was significantly lower than in melanoma and lung cancer (Extended Data Fig.5H). Consistent with their low expression, we could detect PD1-PDL1 protein complex formation via A-FRET in only 12 of the 58 analysed regions (Supplementary Table 2). Moreover, the values of A-FRET intensity were lower than in melanoma and renal cancer^16^. The proportion of regions with detectable PD1-PDL1 complex was significantly less in hypermutated than in non-hypermutated CRCs, while there was no difference between DB- and nDB-CRCs (Fig.4M).

Our analyses show an enrichment of CD74^+^ macrophages in CRCs that respond to immune checkpoint inhibitors. PD1 and PDL1 are overall lowly expressed at the gene and protein levels and show no association with response indicating that, unlike other cancer types^2^, they cannot be used as biomarkers of response to anti-PD1 immunotherapy in CRC.

### CD74^+^PDL1^+^ macrophages interact with PD1^+^ cytotoxic and proliferating CD8 T cells

Our deep investigation of immune infiltrates showed that DB-CRCs are immune hot tumours, with high levels of CD74^+^ macrophages compared to nDB-CRCs as well as of cytotoxic and proliferating T cells associated with their hypermutated phenotype. Because CD74^+^ macrophages also expressed PDL1 while the two populations of CD8 T cells expressed PD1 (Supplementary Table 7, Extended Data Fig.4), we asked whether these cells were proximal in the TME and interacted through PD1-PDL1 contact.

To interrogate this, we identified CD8^+^GzB^+^ and CD8^+^Ki67^+^ cells in the validation cohort (Fig.5A) and in all hypermutated CRCs (Fig.5B) applying a threshold of 0.05 (GzB) and 0.15 (Ki67) expression to all CD8 T cells. We verified that these cells had the same phenotypic features of cluster 1 (CD8^+^GzB^+^ cells) and cluster 2 (CD8^+^Ki67^+^ cells) of the discovery cohort (Extended Data Fig.6A-C). These two subpopulations did not selectively express any additional T cell markers used in the validation cohort, except the immune checkpoint protein LAG3 (Fig.5A). The absence of TCF7 expression in proliferating CD8 T cells suggested that they do not have stem-like characteristics and are not analogous to recently described intratumoural T cell developmental niches^17^,^18^.

**Fig. 5.**
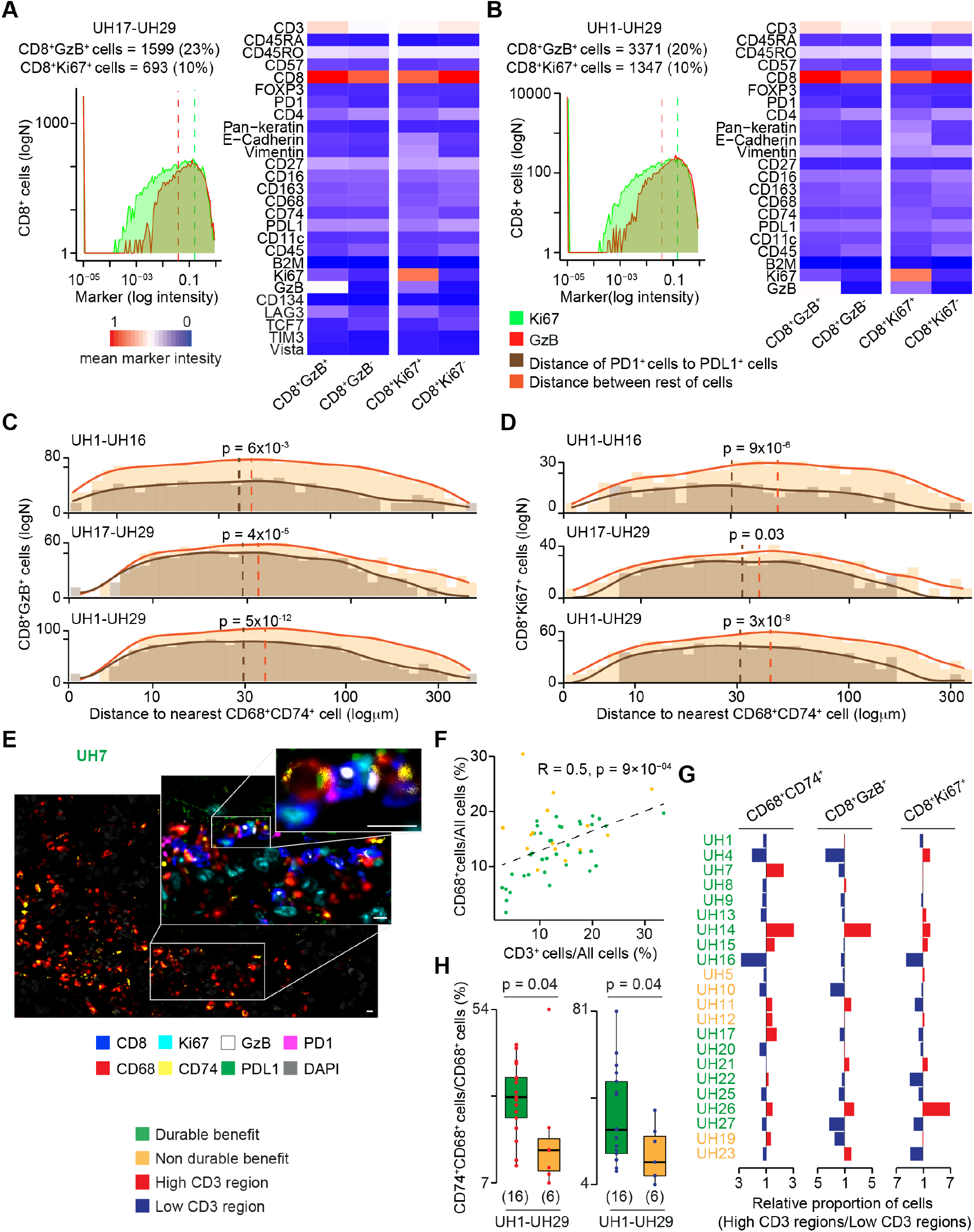
Interaction between CD74^+^ macrophages and GzB^+^Ki67^+^ CD8 T cells. CD8^+^GzB^+^ and CD8^+^Ki67^+^ T cells in the validation (**A**) and combined (**B**) cohorts were identified applying a threshold of 0.05 GzB and 0.15 Ki67 expression to CD8 T cells, respectively following confirmation by IMC image histological inspection. The mean intensities of IMC markers in CD8^+^GzB^+^ or CD8^+^Ki67^+^ and CD8^+^GzB- or CD8^+^Ki67-T cells are reported. Colour scale was normalised across all markers and cells. Distance distributions of CD8^+^GzB^+^ (**C**) or CD8^+^Ki67^+^ (**D**) to the nearest CD74^+^ macrophage in the discovery, validation and combined cohorts. Distances between cells were divided into 1,1μm bins and the density curves fitting the histograms were measured. Distributions of PD1^+^ or PDL1^+^ and rest of cells were compared using two-sided Wilcoxon test. Dashed lines represent medians of the distributions. **(E)** High resolution mIF image of a representative CRC with highlighted a cluster of CD74^+^ macrophages (main image) and their interactions with CD8^+^GzB^+^ and CD8^+^Ki67^+^ T cells (zoom-ins). Image was scanned at 40x magnification. Scale bars = 10μm. **(F)** Correlation between normalised T cells and macrophages in 26 DB- and nDB-CRCs of the discovery and validation cohorts. Pearson correlation coefficient R and associated p-value are shown. **(G)** Ratios of normalised CD74^+^ macrophages, CD8^+^GzB^+^ and CD8^+^Ki67^+^ T cells between regions with high and low T cell infiltration in each tumour. For samples with more than two regions, the total cells in the high or low regions were normalised and used to compute the ratio. **(H)** Comparisons of normalised of CD74^+^ macrophages between DB and nDB-CRCs of the combined cohorts considering only high (left) and low (right) T cell Infiltration regions. Distributions were compared using a two-sided Wilcoxon test. In **(G-H)**, only 22 DB- and nDB-CRCs with at least two regions were considered.

After identifying the CD8^+^GzB^+^ and CD8^+^Ki67^+^ T cell subpopulations, we measured the centroid distance between them and CD68^+^CD74^+^ cells in all regions analysed. We then compared the distance between CD8^+^GzB^+^PD1^+^ or CD8^+^Ki67^+^PD1^+^ and CD68^+^CD74^+^PDL1^+^ cells to the distance between all other cells. We found that CD8^+^GzB^+^PD1^+^ and CD8^+^Ki67^+^PD1^+^ cells were closer to CD68^+^CD74^+^PDL1^+^ cells than other cells in the discovery, validation and combined cohorts (Fig.5C,D). Moreover, a substantial fraction of CD74^+^ macrophages (52% in DB-CRCs and 32% in nDB-CRCs) aggregated in high-density clusters composed of ≥5 cells/10,000*μ*m^217^. These computationally identified clusters of CD74^+^ macrophages were proximal to CD8^+^GzB^+^ and CD8^+^Ki67^+^ cells (Extended Data Fig.7). The existence of these clusters was confirmed through independent histological assessment of IMC images (Extended Data Fig.8). This also allowed visualisation of direct interactions between CD8^+^GzB^+^PD1^+^ or CD8^+^Ki67^+^PD1^+^ and CD68^+^CD74^+^PDL1^+^ cells. To observe these interactions at higher resolution, we performed mIF with eight key markers defining CD74^+^CD68^+^, GzB^+^CD8^+^and Ki67^+^CD8^+^ cells (Supplementary Table 5). We confirmed the presence of clusters of CD74^+^ macrophages in close proximity to CD8^+^GzB^+^ and CD8^+^Ki67^+^ T cells and detected their interaction via PD1-PDL1 contact (Fig.5E, Extended Data Fig.9).

Taking advantage of the multiple profiled regions per tumour, we asked how the observed intra-tumour T cell heterogeneity (Fig.1C, Extended Data Fig.1D) impacted on the distinctive infiltration pattern of DB-CRCs. First, we observed that tumour regions rich in T cells were also rich in macrophages (Fig.5F), indicating that intra-tumour heterogeneity involves a more general pattern of immune co-infiltration. Next, we investigated how the three key populations of CD8^+^GzB^+^, CD8^+^Ki67^+^ and CD68^+^CD74^+^ cells were distributed across regions of the tumour and observed that their relative proportions were variable between high and low infiltrate regions (Fig.5G). Despite the heterogeneous composition of the immune infiltrates, we observed consistently higher proportion of CD74^+^ macrophages in DB-CRCs when compared to nDB-CRCs independently of the T cell infiltration levels (Fig.5H).

Since CD74^+^ macrophages consistently differ between DB and nDB-CRCs across cohorts and regions, we propose that their interaction with CD8^+^GzB^+^PD1^+^ and CD8^+^Ki67^+^PD1^+^ cells through PDL1 is key to confer durable benefit from treatment.

## DISCUSSION

In this study, we profiled multiple CRC regions using a multi-omic and multi-regional approach that allowed the integration of genomic, transcriptomic, histopathologic and immune-phenotypic data to characterise the tumour-immune interactions determining response to immune checkpoint blockade.

After extensive unsupervised investigation of variability in leukocyte subpopulations between DB and nDB-CRCs using multiple approaches, CD74^+^ macrophages were the only immune cell population that consistently segregated with response. Remarkably, despite the widespread genetic and immune inter- and intratumour heterogeneity and independently of whether immunotherapy was given as first line, second line or in the chemorefractory setting, high levels of CD74^+^ macrophages were a consistent feature of DB-CRCs. These macrophages express PDL1 and make direct contact with PD1^+^ CD8 T cells that are enriched in all hypermutated CRCs. Our data suggest that this is a key interaction that restrains CD8 T cell function as cancer progresses, and the one that anti-PD1 antibodies break to release cytotoxic anti-tumour activity.

The high CD8 levels in hypermutated CRCs are likely enabled by the observed low activation of the WNT pathway, resulting in an immune hot environment. To evade immune elimination, responsive hypermutated CRCs develop immune escape mechanisms, via either genetic inactivation or transcriptional repression of antigen presenting genes. Interestingly, unresponsive hypermutated CRCs do not show such a pervasive disruption of the antigen presentation machinery, despite comparably high levels of CD8 infiltration. The molecular mechanisms by which these tumours survive the attack of cytotoxic CD8 T cells need further investigation although a possible cellular explanation could reside in their significantly reduced proportion of CD74^+^ macrophages.

Our study highlights CRC-specific traits of response to anti-PD1 immunotherapy. For example, we showed that in CRC high TMB is necessary but not sufficient to achieve durable benefit and that, above the critical threshold of hypermutated phenotype, even CRCs with very high TMB may not respond to treatment. This is different from lung cancer and melanoma where response always positively correlates with TMB ^7^,^11^. It also suggests that, at least in CRC, a low TMB is a marker of resistance not because of a low neoantigenic load but because it is indicative of a different biology leading to immune cold tumours. Moreover, while the impairment of antigen presentation in immune hot tumours is shared with other cancer types ^19^, the association of *B2M* loss with response and the overall low expression of PD1 and PDL1 are specific to CRC. This suggests that universal predictive markers of response to immunotherapy across tumour types may not exist and that the histo-genetic origin of the tumour as well as the features of the TME should be considered. In the case of CRC, these may include clonal immunogenic mutations and expanded T cells, low activation of the WNT pathway and high infiltration of CD8 T cell s coupled with CD74 macrophages.

## METHODS

### Patient cohorts, treatment and experimental design

FFPE blocks were obtained from the resection of the primary tumour or local relapse of 16 patients (discovery cohort) and 13 patients (validation cohort) treated with immune checkpoint inhibitors in the setting of metastatic CRC until disease progression, unacceptable toxicity or completion of treatment (Supplementary Table 1). In the discovery cohort, patients UH1-UH10 were treated with Pembrolizumab (200 mg every three weeks) as part of the KEYNOTE 177 clinical trial (ClinicalTrials.gov, NCT02563002)^5^. Patients UH11-UH16 were treated with Nivolumab (240mg every two weeks). In the validation cohort, patients UH17-UH19 were part of the KEYNOTE 177 trial, UH26 received Pembrolizumab (2mg/kg every three weeks) and patients UH20-UH25 and UH29 were treated with Nivolumab (240 mg every two weeks). Patient UH27 received Ipilimumab (1mg/kg) in combination with Nivolumab (3mg/kg) every three weeks for four cycles followed by Nivolumab alone (240 mg every two weeks). Patient UH28 received Nivolumab (3mg/kg) for two cycles, then Ipilimumab (1mg/kg) in combination with Nivolumab every three weeks for three cycles. Patients treated with Nivolumab were enrolled in the UK wide Bristol Myers Squibb Individual Patient Supply Request Programme as per Article 5/1 of Article Directive 2001/83/EC. All patients were consented at the UCL Cancer Institute Pathology Biobank - REC reference 15/YH/0311.

Response to therapy was assessed using the formal Response Evaluation Criteria in Solid Tumours (RECIST)^20^ version 1.1. Patients were considered to have durable benefit (DB) from the treatment if the disease did not progress for at least 12 months after commencing immunotherapy, except for patient UH28 who showed partial response for >10 months at the time of writing. Patients were considered to have no durable benefit (nDB) from treatment if the disease progressed within 12 months. Twelve-month cut-off was considered clinically better than the progression-free survival from chemotherapy as first line treatment of metastatic stable (8.3 months^21^) or hypermutated (8.2 months^5^) CRC.

Twenty-four sequential sections were cut from each FFPE tumour block of samples UH1-UH16 with a microtome. Sections were then used for CD3 staining (slides A, B, F, H and J); Imaging Mass Cytometry (IMC, slide C); multiplexed Immunofluorescence (mIF, slide D); Whole Exome Sequencing (WES, slides E1-5); RNA sequencing (RNA-seq, slides G1-5); T-Cell Receptor Beta sequencing (TCR-seq, Slides I1-5) and to detect PD1-PDL1 interaction *in situ* (slides K1-2). For samples UH17-UH23 and UH25-UH27, 11 sequential sections were used for CD3 staining (slides A, E and G); Haematoxylin and Eosin (HE) staining (slide B); IMC (slide C); mIF (slide D) and RNAseq (slides F1-5). Since UH24, UH28 and UH29 were biopsies, RNAseq was not performed because the tissue was insufficient, and four sequential sections were cut and used for CD3 and HE staining (slides A and B); IMC (slide C) and mIF (Slide D). Sections used for CD3 and HE staining, IMC, mIF and A-FRET were 4μm thick, while those used for DNA and RNA extraction were 10μm thick. Tumour content was assessed by a board-certified surgical pathologist (M.R.J.).

### CD3 staining and quantification

CD3 staining was performed upon slide dewaxing and heat-induced epitope retrieval (HIER) using Antigen Retrieval Reagent-Basic (R&D Systems). Tissues were blocked and incubated first with anti CD3 antibody (Dako, Supplementary Table 5) and subsequently with horseradish peroxidase conjugated anti rabbit antibody (Dako). They were then stained with 3,3’ diaminobenzidine (DAB) substrate (Abcam) and haematoxylin. Slide A was reviewed by a certified pathologist (M.R-J) to identify two to four regions per slide with high and low CD3^+^ infiltrating cells (for a total of 90 regions, Supplementary Table 2), preferably in proximity to the invasive margins of the tumour (Extended Data Fig.1B).

Digital acquisition of CD3 stained slides was performed using Hamamatsu Nanozoomer (Hamamatsu Photonics) or Axioscan Z1 (Zeiss) at 20x resolution. The whole slide images were then loaded into QuPath^22^ v0.2.0-m4 to quantify CD3^+^ infiltration within each selected region. The “Estimate Stain Vector” function was run as pre-processing step to increase the contrast between DAB and haematoxylin stains. The outlines of the regions delimited by the pathologist in slide A were projected in all other slides (Extended Data Fig.1C). The regions were then divided into 0.09 mm^2^ large tiles and CD3^+^ cells were quantified within each tile using the “positive cell detection” function that first detects all the cells present in the tile and then identifies positive cells based on the DAB intensity threshold. The median value of CD3^+^ cells per mm^2^ across all tiles was considered as representative of CD3^+^ infiltration for that region. For slides B and I, CD3^+^ cells were also quantified for the whole tumour region.

### Imaging mass Cytometry (IMC)

Two panels of 35 antibodies in total were assembled to represent the main immune and stromal populations of the gut TME (panels I and II). Twenty-five of these antibodies were already metal-tagged (Fluidigm), while ten were purchased in a carrier-free form, tested via immunohistochemistry in FFPE appendix sections and tagged using the Maxpar X8 metal conjugation kit (Fluidigm). To identify the optimal dilution for each antibody, concentrations ranging from 1/50 to 1/5,000 were tested in FFPE appendix sections. After staining and ablation, images were visualised using MCD Viewer (Fluidigm) and the concentration giving the best signal to background ratio was selected (Supplementary Table 5).

IMC was performed in 38 regions of the discovery cohort using panel I and in 22 regions of the validation cohort using panel II (Supplementary Table 2). In the discovery cohort, the two regions (one with low and one with high CD3 infiltration) with the highest tumour content were selected except for UH4 UH6, UH9 and UH12. For UH6, UH9 and UH12, all four regions were analysed, for UH4, the two high and two low CD3 regions were analysed together, to be consistent with WES and RNA-seq analyses (see below). In the validation cohort, the two regions with the highest difference in CD3 infiltration were selected except for the three biopsies (UH24, UH28 and UH29) and UH18, where only one region was analysed. Slides were incubated for one hour at 60°C, dewaxed, rehydrated and subjected to HIER using a pressure cooker and Antigen Retrieval Reagent-Basic (R&D Systems). Tissues were blocked in a solution containing 10% BSA (Sigma), 0,1% Tween (Sigma), 1:50 Kiovig (Shire Pharmaceuticals) Superblock Blocking Buffer (Thermo Fisher) for two hours at room temperature. The primary antibody mix was prepared in blocking solution at the selected concentration for each antibody and incubated overnight at 4°C. Slides were then washed twice in PBS and PBS-0.1% Tween and incubated with 2 isotopes (^191^Ir and ^193^Ir) of DNA intercalator Cell-ID™ Intercalator-Ir (Fluidigm) 1.25mM diluted in PBS for 30 minutes at room temperature. Slides were then washed once in PBS and once in MilliQ water and air-dried. Stained slides were loaded in the Hyperion Imaging System (Fluidigm) imaging module to obtain light-contrast high resolution images of approximately 4 mm^2^. For each region, a 1 mm^2^ area with high tumour content and representative of the median CD3^+^ content of the region was selected for laser ablation (Extended Data Fig.3A) at 1 μm/pixel resolution and 200Hz frequency.

### IMC pixel analysis

The main steps of IMC data analysis are summarised in Fig.3A. For each of the 60 ablated regions, TIFF images from each antibody and two DNA intercalators were obtained from the raw IMC .mcd and .txt files using imctools^23^. Pixel intensities for each channel were normalised to the 99^th^ percentile of the intensity distribution and the obtained values scaled between 0 and 1. Background pixels were removed using global thresholding with CellProfiler^24^ 3.1.8. After visual inspection, channels for PD1, PDL1, GzB, CD45RA in both cohorts and TIM3, Vista, TCF7 and CD134 in the validation cohort, were further filtered using probability masks for background and non-background pixels produced with Ilastick^25^ 1.3.0. For this purpose, random forest classifiers were trained using closely related markers (CD3 for PD1; Vimentin for PDL1; CD8 and CD15 for GZB; CD45 and CD45RO for CD45RA). The resulting background probability masks were converted into binary images with CellProfiler^24^ 3.1.8 and applied to the original normalised images to remove the background. Custom R scripts were used to count the positive pixels in all processed images for each channel. The sum of all positive pixels for a channel constitutes the positive area for that channel. Given B2M low expression, an ad-hoc threshold was applied on the normalized intensity and all pixels higher than 0.5 were considered as positive. For regions UH19_87, UH27_96, and UH27_97, the CD3 masking threshold was adjusted after manual inspection to 0.175, 0.15 and 0.15 respectively.

Tumour masks were generated with CellProfiler^24^ 3.1.8 summing up the Pankeratin and E-cadherin channels for all regions except UH18_103 and UH22_112, where only E-cadherin was used. The resulting images were smoothened with a Gaussian filter and filling up all <30 pixel negative areas. For the discovery cohort, the stroma masks were obtained with same procedure using the Vimentin, SMA and DNA channels as input. For the validation cohort, the stroma masks were obtained by summing up DNA, Vimentin, CD68, CD11c, CD3, CD27 and CD45 channels and filling up all the <20 pixel negative areas. The tissue mask for each region corresponded to the sum of tumour and stroma masks. Pixel analysis was performed by normalising the positive areas for each marker or combination of markers over the total tissue area or the area of the five main immune populations (T cells, B cells, macrophages, dendritic cells and neutrophils) for the discovery cohort and for T cell and macrophages only for the validation cohort (Supplementary Table 6).

### IMC single cell analysis

Single cell analysis was based on cell segmentation, assignment of cell identity and phenotype clustering (Fig.3A).

Cell segmentation was performed with CellProfiler^24^ 3.1.8 identifying nucleus and membrane of each cell in each region. First, the two DNA channels were multiplied and used for nucleus segmentation using local Otsu thresholding. Second, all channels for the membrane markers (CD3, CD20, CD27, CD16, CD11c, CD15, SMA, CD34, Vimentin and Pan-keratin for UH1-UH16 and CD45, Pan-keratin and E-cadherin for UH17-UH29) were used to obtain membrane images. Cell masks were then generated by radially expanding each nucleus up to 10 pixels on the membrane mask and only cells overlapping with the tissue mask were retained. Finally, the mean intensity of all markers was measured in each cell in each region.

Cell identities were assigned according to the maximum overlap of the cell area with marker-specific thresholds identified by the histologist (J.S.) after image manual inspection. These thresholds were: ≥25% of CD3^+^ mask for T cells; ≥10% of CD11c^+^ CD68-mask for dendritic cells; >10% of the sum of CD68^+^ CD11c^+^ and CD68^+^ CD11c-masks for macrophages; ≥5% of IgA^+^, IgM^+^, CD20^+^, and CD27^+^ mask for B cells; and ≥25% of CD15^+^ mask for neutrophils. Cells that did not overlap with any of these markers were defined as tumour cells if they overlapped ≥80% with the tumour mask or were left unassigned otherwise. Within CD3^+^ cells, PD1^+^ cells were identified as those showing ≥1% overlap with the PD1 mask. PDL1^+^ cells were identified as those overlapping ≥10% of the PDL1 mask.

For the discovery cohort, single cell phenotype clustering was performed for T cells, B cells, macrophages, dendritic cells, neutrophils, PD1^+^ and PDL1^+^ cells separately using Seurat^26^ 2.4 with custom R scripts for IMC data analysis. Independent clustering was used to compare the relative abundance of cell subpopulations between hypermutated and non-hypermutated CRCs or DB- and nDB-CRCs using Pembrolizumab and Nivolumab samples alone or combined. The total number of cells used in each clustering is shown in Supplementary Table 9. For each main population, the clustering was based on the mean expression of a set of markers typical of that population (Supplementary Table 5). The mean marker intensities across all cells were integrated using multiple canonical correlation analysis (CCA) and aligning the CCA subspaces to reduce the inter-sample variability. The resulting CCA vectors were then used as input for unsupervised clustering with ten values of resolution, ranging from 0.1 to 1.0. The ten resulting sets of clusters were manually inspected and the one with the highest number of biologically meaningful clusters was chosen. For the validation cohort, T cells, macrophages, PD1^+^ and PDL1^+^ cells were identified as described above. CD74^+^ macrophages and CD8^+^GzB^+^ CD8^+^Ki67^+^ T cells were identified using specific expression thresholds on the mean cell intensity (0.1 for CD74; 0.1 for CD8; 0.05 for GzB; 0.15 for Ki67). CD8^+^ T cells positive for both the Ki67 and the GzB threshold were identified as CD8^+^GzB^+^ or CD8^+^Ki67^+^ T cells according the marker with the highest intensity. T cells in the validation cohort underwent single cell clustering, using all 17 T cell markers (Supplementary Table 5). The distribution of cells within each cluster over the total cells was compared between DB- and nDB-CRCs or hypermutated and non-hypermutated CRCs using two-sided Wilcoxon test, correcting for FDR.

### IMC neighbour and cluster density analysis

The pixel coordinates of the centroid of each cell were extracted from the cell masks with CellProfiler^24^ 3.1.8 and used to measure the Euclidean distances between each pair of cells in each region. High-density clusters of CD68^+^CD74^+^ within each region were identified using DBSCAN (Density-Based Spatial Clustering of Applications with Noise^27^) as implemented in the fpc R package version 2.2.5. Starting form cell pixel coordinates, highly dense clusters were defined as portions of the ablated regions with ≥5 CD68^+^CD74^+^per 10,000μm^2^, corresponding to a minimum number of five points (MinPts) within a radius (eps) of 56.42μm.

### Multiplexed Immunofluorescence (mIF)

mIF was performed on slide D of 24 DB- and nDB samples (Supplementary Table 2). An automated Opal-based mIF staining protocol was developed using a Ventana Discovery Ultra automated staining platform (Roche) with eight markers specific for CD8^+^GzB^+^PD1^+^, CD8^+^Ki67^+^PD1^+^ and CD68^+^CD74^+^PDL1^+^ cells, DAPI and Opal fluorophores (Supplementary Table 5). Antibody dilution, incubation time and effect of denaturation steps as well as Opal dilution were assessed for each marker following manufacturer’s instructions. The optimal antibody-Opal pairing was achieved considering the expected expression and cellular localisation of each marker and the fluorophore brightness to minimize fluorescence spillage. The final staining order was CD74, TCF7, PDL1, Ki67, PD1, GzB, CD68 and CD8.

Slides were baked for 1 hr at 60°C, loaded onto the autostainer and subjected to a fully automated staining protocol involving deparaffinisation (EZ-Prep solution, Roche), HIER (DISC. CC1 solution, Roche) and seven sequential rounds of 1 hr incubation with the primary antibody, 12 minutes incubation with the HRP-conjugated secondary antibody (DISC. Omnimap anti-Ms HRP RUO or DISC. Omnimap anti-Rb HRP RUO, Roche) and 16 minute incubation with the Opal reactive fluorophore (Akoya Biosciences). For the last round of staining, tissues were incubated with Opal TSA-DIG reagent (Akoya Biosciences) for 12 minutes and with Opal 780 reactive fluorophore for 1 hour (Akoya Biosciences). Before each round of staining, a denaturation step (100°C for 8 minutes) was introduced to remove the primary and secondary antibodies from the previous cycle without disrupting the fluorescent signal. Once the staining was completed, the slides were counterstained with 4’,6-diamidino-2-phenylindole (DAPI, Akoya Biosciences) and coverslipped using ProLong Gold antifade mounting media (Thermo Fisher Scientific). Fluorescently labelled slides were scanned using a Vectra Polaris automated quantitative pathology imaging system (Akoya Biosciences). Spectral libraries were constructed with the inForm 2.4 image analysis software (Akoya Biosciences) following the manufacturer’s instructions. Whole-slide scans were obtained at 20x and 40x magnification using appropriate exposure times, and several fields of views were selected per slide and loaded into inForm (Akoya Biosciences) for spectral unmixing and autofluorescence isolation using the spectral libraries.

### DNA sequencing

All regions were macro-dissected with a needle under a stereo microscope using slide A as a guide (Extended Data Fig.1E). Genomic DNA was extracted from 32 tumour regions and 16 matched normal tissue of slides E1-5 of samples UH1-UH16 (Supplementary Table 2) and the two regions corresponding to those used for IMC were selected. For UH4, the two high and low CD3^+^ regions were merged to obtain enough DNA for library preparation. DNA was extracted using GeneRead DNA FFPE kit (Qiagen) and DNA libraries were prepared using 50-200ng of genomic DNA with the KAPA HyperPrep kit (Roche). Protein-coding genes were captured using SureSelectXT Human All Exon V5 probes (Agilent) and sequenced on Illumina HiSeq 4000 using 100bp paired end read protocol, according to manufacturer’s instructions. Approximately 100 million reads were generated per sample.

Raw reads were aligned to GRCh38 reference human genome using BWA MEM^28^ v0.7.15 after pre-alignment quality control. Regions harbouring small insertions and deletions (indels) were re-aligned locally using GATK^29^ v3.6. The resulting BAM files were sorted, merged, marked for duplicates and subjected to post-alignment quality control using Picard v2.10.1. The final mean depth of coverage was >70x for tumour and >30x for normal samples, considering only targeted exons as defined in the SureSelectXT BED file (50.5Mbp in total).

Somatic SNVs and indels were called using Strelka^30^ v2.9.0 on the targeted exome extended 100bp in both directions. Mutations were retained if they had an Empirical Variant Scoring (EVS) >7 for SNVs and >6 for indels in at least one region of the same patient. Mean sensitivity in variant calling was >91% in all patients expect UH16 (33%) as assessed using 241 somatic mutations from FM1^31^ or the patient pathological reports for comparison. Nineteen mutations in FM1 or pathological reports but missed by Strelka were added to the pool of somatic alterations after manual check.

For samples UH1-UH3, UH7-UH10 and UH12, copy number analysis was done using ASCAT^32^ v2.5.2. To process WES data, AlleleCount^33^ v4.0.0 was run on germline SNPs from 1000 Genomes Phase 3^34^ after correction for GC bias. For each SNP, a custom script was used to calculate the LogR and B-Allele Frequency (BAF). SNPs with <6 reads were filtered out in all samples except UH1 and UH10 where 5 or 7 reads were used. Because of degraded starting DNA of samples UH4-UH6, UH11 and UH13-UH16, the DepthOfCoverage option of GATK^29^ v3.6 was used to calculate SNP LogR and copy numbers for genomic segments were obtained using Copynumber R package. The gene copy number was derived from that of the genomic segment covering at least 25% of the gene length.

### Prediction of damaged genes and immunogenic mutations

ANNOVAR^35^ (release 16/04/2018) was used to annotate exonic or splicing SNVs and indels. All truncating mutations (stop-gains, stop-losses and frameshift indels) were considered as damaging. Non-truncating mutations (non-frameshift indels and missense SNVs) were considered damaging if predicted by at least five function-based methods or two conservation-based methods^36^. Mutations within two bps of a splicing junction were considered as damaging if predicted by at least one ensemble algorithm in dbNSFP. Gain of function mutations were predicted using OncodriveClust^37^ for the Pembrolizumab and the Nivolumab patients together, with default parameters and with false discovery rate (FDR) <10%.

A gene was considered amplified if its copy number was >1.4 times the sample ploidy or >2 if the ploidy was not available in both regions of the same patient, or if its CPM was >1.5 than in the other region of the same patient. A gene was considered as deleted if it had copy number = 0, CPM = 0 and had no mutations. A gene was considered deleted in heterozygosity if it had copy number = 1.

Amplified genes, deleted genes, heterozygously deleted genes with a damaging mutation in the other allele and copy number neutral genes with at least one damaging mutation were considered as damaged genes.

To predict putative immunogenic mutations from all somatic SNVs and indels, HLA typing of each patient was predicted from the normal BAM files using Polysolver^38^ v4 (Supplementary Table 9). NeoPredPipe^39^ was then used to predict neoantigens in expressed genes (CPM >0), with a strong HLA binding (rank <0.5%) and crossreferenced with known epitopes (UniProt reference proteome). SNVs or indels generating at least one neoantigen were considered as potentially immunogenic. The neoantigenic index was calculated for each region as:

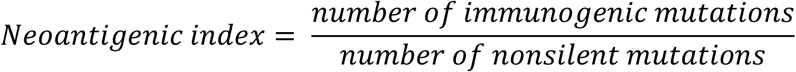

PyClone^40^ v.0.13.1 was run to assess the clonality of predicted immunogenic mutations, defined as the proportion of tumour cells harbouring the mutation. PyClone was run independently for each region using tumour purity from the pathological assessment of slide A (Supplementary Table 2) and the gene copy number from ASCAT.

### RNA sequencing

Total RNA was extracted from 58 macro-dissected regions of slides G1-5 in samples UH1-UH16 and 30 regions of slides F1-5 in samples UH17-UH23 and UH25-UH27 (Supplementary Table 2) using the High Pure FFPE RNA isolation kit (Roche). For UH4, the two high and low CD3^+^ regions were merged to obtain enough RNA for library preparation. RNA libraries were prepared starting from 5-50 ng of RNA using the QuantSeq 3’mRNA-seq Library Prep kit FWD for Illumina (Lexogen) and sequenced on Illumina HiSeq 4000 using 75 or 100 bp single end reads, according to manufacturer’s instructions. Approximately 5-40 million reads were generated per sample.

Raw reads were processed using the Lexogen QuantSeq 3’ mRNA-seq pipeline with default parameters^41^. Reads were first trimmed to remove Illumina adapters and polyA tails using bbduk from BBMap^42^ v36.20. Trimmed reads were then aligned to GRCh38 reference human genome using STAR^43^ v2.5.2a. Between 50-98% of the initial reads were retained after alignment and quality check. Gene expression was quantified using HTSeq^44^ v0.6.1p1 and the GDC h38 GENCODE v22 GTF annotation file. To account for differences in sequencing depth across regions raw counts were normalised to the counts-per-million (CPM) gene expression unit calculated as:

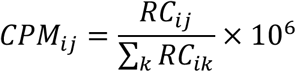

where *RC_ij_* is the raw read count of gene (*j*) in region (*i*). Since for samples UH1-UH16 RNA-seq was performed in six batches, potential batch effect was corrected using removeBatchEffect function from the Limma package^45^ v3.36.5 on the log2-transformed CPM matrix with default parameters.

Differential gene expression between tumour groups was assessed using DESeq2^46^ v1.20.0 from the raw read counts with default parameters with alpha set to 5%. To account for the experimental and clinical variability across samples (Supplementary Table 1), uncorrelated batch and clinical co-variates were included in the analysis (discovery cohort: batch effect, prior lines of treatment and Lynch syndrome in the comparison of DB- and nDB-CRCs of the; batch effect in the comparison of hypermutated and non-hypermutated CRCs. Validation cohort: prior lines of treatment, Lynch syndrome, treatment type and TNM staging).

A gene was considered as differentially expressed if DESEq2 Wald test FDR was <5% and had a fold change |>2|. Differentially expressed genes were used for pathway enrichment analysis in the three comparisons using MetaCore v20.3 build 70200 (Clarivate Analytics). Pathway enrichment was assessed through over-representation analysis based on a hypergeometric test. A pathway was considered enriched if the FDR was <10%.

### TCR sequencing

TCR-seq was performed in 28 macro-dissected regions in slides I1-5 of the discovery cohort, after excluding UH1, UH4 and UH5 because DNA was not sufficient (Supplementary Table 2). For UH12 all four regions were sequenced while for UH16 the two high and low CD3^+^ regions were merged. For all remaining samples, the two regions corresponding to those used for IMC and WES were used.

DNA was extracted from macro-dissected regions (Extended Data Fig.1E) using GeneRead DNA FFPE kit (Qiagen) and submitted to Adaptive Biotechnologies (Seattle, USA) for non-lymphoid tissue (survey level) TCR-seq using a two-step, amplification bias-controlled multiplex PCR approach^47^. In the first step, V and J gene segments encoding the TCR beta CDR3 locus were amplified using reference gene primers to quantify total nucleated cells and measure the fraction of T cells in each sample. In the second step, proprietary barcodes and Illumina adapters were added. Finally, CDR3 and reference gene libraries were sequenced according to the manufacturer’s instructions.

Raw reads were de-multiplexed and processed to remove adapter and primer sequences, identify and remove primer dimer, germline and other contaminant sequences. Resulting reads were clustered using both the relative frequency ratio between similar clones and a modified nearest-neighbour algorithm, to merge closely related sequences to correct for technical errors introduced through PCR and sequencing. The resulting reads were sufficient for annotating the V(N)D(N)J genes of each unique CDR3 and the translation of the encoded CDR3 amino acid sequence. V, D and J gene definitions were based on annotation in accordance with the IMGT database (www.imgt.org). The set of observed biological TCR Beta CDR3 sequences were normalised to correct for residual multiplex PCR amplification bias and quantified against a set of synthetic TCR Beta CDR3 sequence analogues^47^. Data was analysed using the immunoSEQ Analyzer toolset.

### Detection of PD1-PDL1 interaction *in situ*

A total of 58 regions in slides K1-2 of the discovery cohort (Supplementary Table 2) were submitted to FASTBASE Solutions (Derio, Spain) to measure the interaction between PD1 and PDL1 *in situ* via amplified immune Förster Resonance Energy Transfer (i-FRET)^16^. Slides K1 were incubated overnight at 4 °C with anti PD1 primary antibody (Supplementary Table 5) for donor only analyses. Slides K2 were stained with both anti PD1 and anti PDL1 primary antibodies for donor and acceptor analyses. Slides K1 were subsequently incubated with anti-mouse Fab-ATTO488 and slides K2 with both anti-mouse Fab-ATTO488 and anti-rabbit Fab-HRP. Alexa594 conjugated tyramide was added to slides K1 and K2 at 1/100 dilution in presence of 0,15% H2O2 and incubated at room temperature in the dark for 20 minutes. After washing in PBS and PBST twice, slides were mounted using Prolong Diamond Antifade Mount (Thermo Fisher), sealed and incubated at room temperature overnight before being transferred to a 4 °C refrigerator for storage. FASTBASE Solutions SL frequency domain FLIM automated software programme was used to measure the excited-state lifetime of donor fluorophore (ATTO488) in both K1 and K2 slides. A-FRET efficiency (E) was calculated as:

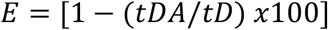

where *tDA* is donor lifetime in Slide K1 and *tD* is donor lifetime in Slide K2. *tDA* and *tD* values were collected for 793 optical fields of view (FOVs, with a median of 12 FOVs per region) in total to cover the whole surface of the regions analysed. Data were then collected in .csv files and imported into a macro spreadsheet programmed to calculate the i-FRET efficiency in each FOV. The results were finally expressed as the median FOV values per region.

## ACKNOWLEDGMENTS

We thank Prof Rebecca Fitzgerald (University of Cambridge) and Prof Toby Lawrence (King’s College London) for critical comments to the manuscript. We acknowledge technical support from the Advanced Sequencing, the Flow Cytometry and the Experimental Histopathology Platforms of the Francis Crick Institute, the National Institute for Health Research Biomedical Research Centre Immune Monitoring Core Facility Centre based at Guy’s and St Thomas’ NHS Foundation Trust and King’s College London, Christopher Applebee and Sánchez-Magraner for the FRET analysis and Joe Brock for graphical illustrations. The views expressed are those of the authors and not necessarily those of the NHS, the NIHR or the UK Department of Health. The results published here are in part based upon data generated by The Cancer Genome Atlas managed by the NCI and NHGRI.

## AUTHOR CONTRIBUTIONS

M.B. developed the computational method and performed IMC data analysis; M.R.K analysed RNA-seq and TCR-seq data and helped with IMC analysis; L.M. performed all immunostaining and analysed the images; A.A.S. performed the mIF experiments with the assistance of P.W., J.K., T.D. and E.N.; A.A.S. and L.B. macro-dissected the regions and prepared samples for genomic and transcriptomic analyses; S.C. and D.T. analysed the WES data; N.P. and K.T. provided support for IMC; R.G., S.W., G.A.W., M.A.B and C.S. provided the protocol for FFPE WES; S.J. assisted RNA-seq analysis; J.M., B.L. and P.P. performed the A-FRET experiments and contributed their interpretation. V. K., E.F., H.T.A. and K.K.S. identified the patients and provided clinical assessments; M.R.J. performed pathological assessments; F.D.C. conceived and directed the study and analysed the data with support of J.S.; F.D.C., M.B., M.R.K., L.M., D.T., S.C., A.A.S., L.B, M.R.J., K.K.S. and J.S. wrote and edited the manuscript. All authors read and approved the final manuscript.

